# Antibody responses to Zika virus proteins in pregnant and non-pregnant macaques

**DOI:** 10.1101/352880

**Authors:** Anna S. Heffron, Emma L. Mohr, David Baker, Amelia K. Haj, Connor R. Buechler, Adam Bailey, Dawn M. Dudley, Christina M. Newman, Mariel S. Mohns, Michelle Koenig, Meghan E. Breitbach, Mustafa Rasheed, Laurel M. Stewart, Jens Eickhoff, Richard S. Pinapati, Erica Beckman, Hanying Li, Jigar Patel, John C. Tan, David H. O’Connor

**Affiliations:** Department of Pathology and Laboratory Medicine, University of Wisconsin-Madison, Madison, WI, United States of America; Department of Pediatrics, School of Medicine and Public Health, University of Wisconsin-Madison, Madison, WI, United States of America; Department of Biostatistics & Medical Informatics, University of Wisconsin-Madison; Technology Innovation, Roche Sequencing Solutions, Madison, WI, United States of America

## Abstract

The specificity of the antibody response against Zika virus (ZIKV) is not well-characterized. This is due, in part, to the antigenic similarity between ZIKV and closely related dengue virus (DENV) serotypes. Since these and other similar viruses co-circulate, are spread by the same mosquito species, and can cause similar acute clinical syndromes, it is difficult to disentangle ZIKV-specific antibody responses from responses to closely-related arboviruses in humans. Here we use high-density peptide microarrays to profile anti-ZIKV antibody reactivity in pregnant and non-pregnant macaque monkeys with known exposure histories and compare these results to reactivity following DENV infection. We also compare cross-reactive binding of ZIKV-immune sera to the full proteomes of 28 arboviruses. We independently confirm a purported ZIKV-specific IgG antibody response targeting ZIKV nonstructural protein 2B (NS2B) that was recently reported in ZIKV-infected people and we show that antibody reactivity in pregnant animals can be detected as late as 127 days post-infection (dpi). However, we also show that these responses wane over time, sometimes rapidly, and in one case the response was elicited following DENV infection in a previously ZIKV-exposed animal. These results suggest epidemiologic studies assessing seroprevalence of ZIKV immunity using linear epitope-based strategies will remain challenging to interpret due to susceptibility to false positive results. However, the method used here demonstrates the potential for rapid profiling of proteome-wide antibody responses to a myriad of neglected diseases simultaneously and may be especially useful for distinguishing antibody reactivity among closely related pathogens.

**Author summary:** ZIKV has emerged as a vector-borne pathogen capable of causing serious illness in infected adults and congenital birth defects. The vulnerability of communities to future ZIKV outbreaks will depend, in part, on the prevalence and longevity of protective immunity, thought to be mediated principally by antibodies. We currently lack diagnostic assays able to differentiate ZIKV-specific antibodies from antibodies produced following infection with closely related DENV, and we do not know how long anti-ZIKV responses are detectable. Here we profile antibodies recognizing linear epitopes throughout the entire ZIKV polyprotein, and we profile cross-reactivity with the proteomes of other co-endemic arboviruses. We show that while ZIKV-specific antibody binding can be detected, these responses are generally weak and ephemeral, and false positives may arise through DENV infection. This may complicate efforts to discern ZIKV infection and to determine ZIKV seroprevalence using linear epitope-based assays. The method used in this study, however, has promise as a tool for profiling antibody responses for a broad array of neglected tropical diseases and other pathogens and in distinguishing serology of closely-related viruses.

## Introduction

Serologic assays designed to detect Zika virus (ZIKV) infection suffer from cross-reactivity with antibodies to closely related dengue virus (DENV), due to the high level of amino acid sequence identity (average ~55%) and structural similarity [1–4] between the two viruses. Humoral cross-reactivity with other similar arboviruses has been reported as well [1–2]. Serologic assays have been developed to detect past ZIKV infection, reporting sensitivity varying from 37% to 97% and specificity varying from 20% to 90% [5–8]. Most of these assays detect antibodies to the ZIKV envelope protein or nonstructural protein 1 (NS1) [7,9–11]. A recent publication by Mishra et al. employed a high-density peptide microarray to identify antibodies to linear ZIKV epitopes lacking cross-reactivity with other flaviviruses [12]. This group identified an IgG immunoreactive peptide sequence in the ZIKV nonstructural protein 2B (NS2B) which induced little antibody binding in serum from ZIKV-naive people and was bound in early convalescence in most cases of symptomatic ZIKV infection [12]. This group did note seropositivity in a ZIKV-naive, DENV-immune individual and in one individual with no known flavivirus infection history.

Because ZIKV, DENV, and other arboviruses are similar in structure and acute clinical syndrome, are spread by the same mosquito vectors [13], and are co-endemic [14–15], it is difficult to identify people who have unequivocally been exposed to ZIKV only, adding uncertainty to efforts to profile ZIKV-specific antibody responses in humans. In contrast, macaques raised in indoor colonies can be infected specifically with ZIKV, DENV, or other pathogens. Macaque models of ZIKV infection provide a close approximation of human ZIKV infection in regards to natural history [16–19], tissue tropism [16-17,20-21], and transmission [17,22–25]. Importantly, macaques infected with ZIKV during pregnancy provide insight into the pathogenesis of congenital ZIKV infection [17-18,21-22,26-29]. Since the strain, dose, and timing of macaque model ZIKV infection is exactly known, the kinetics and specificity of humoral immune responses can be profiled in macaques with better resolution than is possible in cross-sectional human studies.

The peptide microarray technology we use in this study allows for one serum sample to be assayed against six million unique 16-residue peptides, or for 12 samples to each be assayed against 392,000 peptides, on a single chip (Roche Sequencing Solutions, Madison, WI). The technology has been used in proteome-wide epitope mapping [30], profiling of antibody responses in autoimmune disease [31], profiling venom toxin epitopes [32], determining functions of cellular enzymes [33], *de novo* binding sequence discovery [34], epitope validation following phage display screening [35], and screening for tick-borne disease seroprevalence [36]. We previously used this tool to examine antibody responses in simian pegivirus (SPgV) and simian immunodeficiency virus (SIV) infections [37]. As mentioned above, a recent publication explored use of this technology in profiling human antibody responses against flaviviruses [12]. This linear peptide microarray technology has advantages over previous assays through its capacity to screen for reactivity to a large number of pathogens while simultaneously mapping reactive epitopes precisely, and it has the distinct advantage of allowing detection of unexpected epitopes due to its capacity to assay the entirety of a virus’s proteome.

Here, we used high-density peptide microarrays to map macaque IgG epitopes in full-length ZIKV and DENV polyproteins and to compare cross-reactivity to 27 other arboviruses. Our study takes advantage of this technology’s capacity to assess binding to linear epitopes throughout a virus’ entire proteome to identify an epitope in ZIKV NS2B, a protein made intracellularly which embeds in the host cell’s endoplasmic reticulum and thus would not be expected to induce strong antibody responses. We also demonstrate the potential of this technology, through its ability to survey binding throughout many whole viral proteomes simultaneously, to differentiate seroreactivity to an infecting virus from cross-reactivity against a great variety of other similar viruses. These unique aspects of this recently developed peptide microarray technology highlight its capacity to efficiently screen for and identify previously unknown epitopes in a large number of neglected disease-causing pathogens simultaneously and to distinguish infection histories of NTDs, potentially impacting the development of future diagnostics and vaccines.

## Methods

### Macaque study design

Animal demographics, inoculation strain, dose, and route, serum sample collection timelines, and array design used for each animal are described in Table 1. Gestational details and outcomes for pregnant animals are described in Table 2. Additional details on the study histories of the animals in this study can be found at https://go.wisc.edu/b726s1.

**Table 1.**
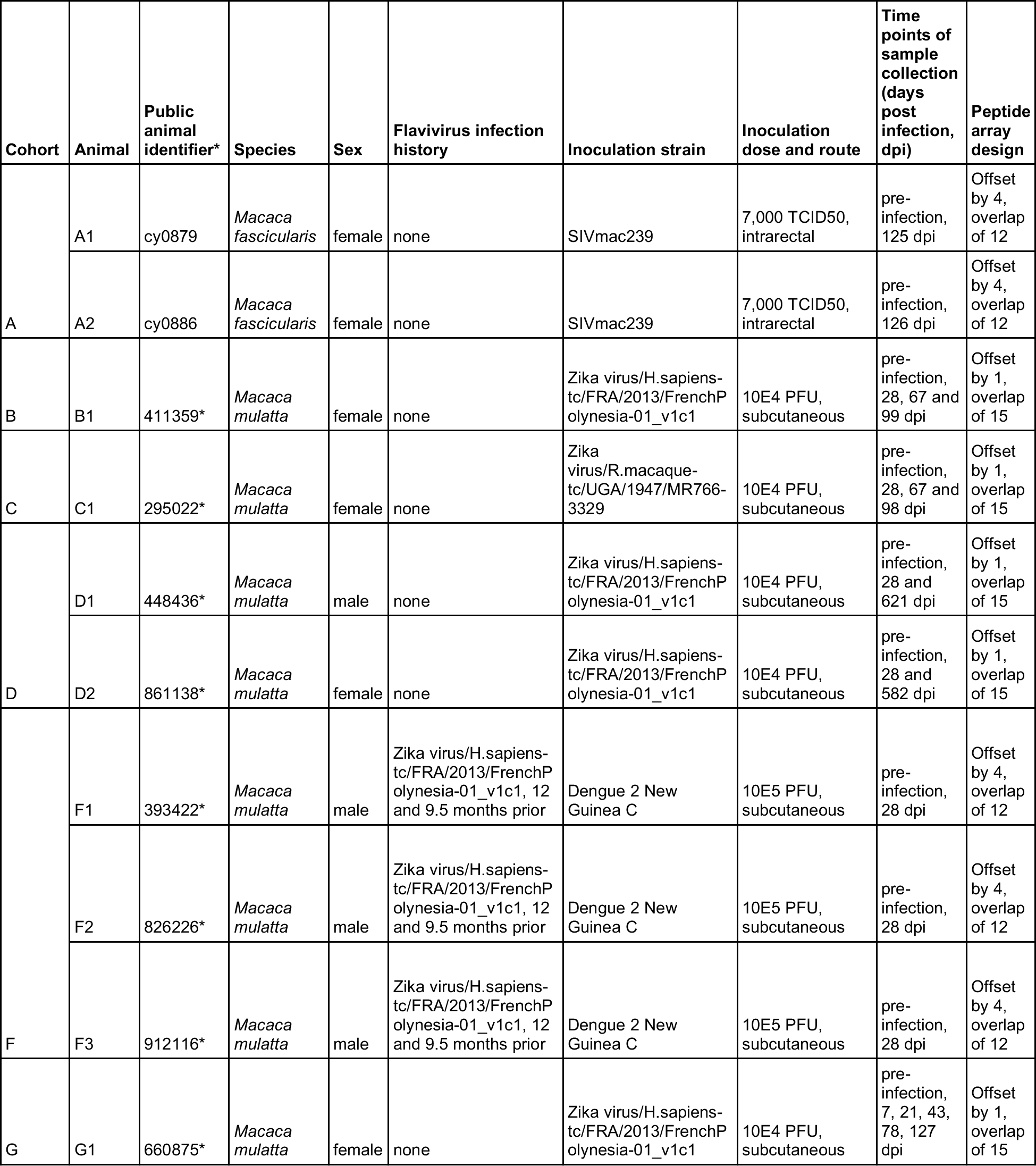
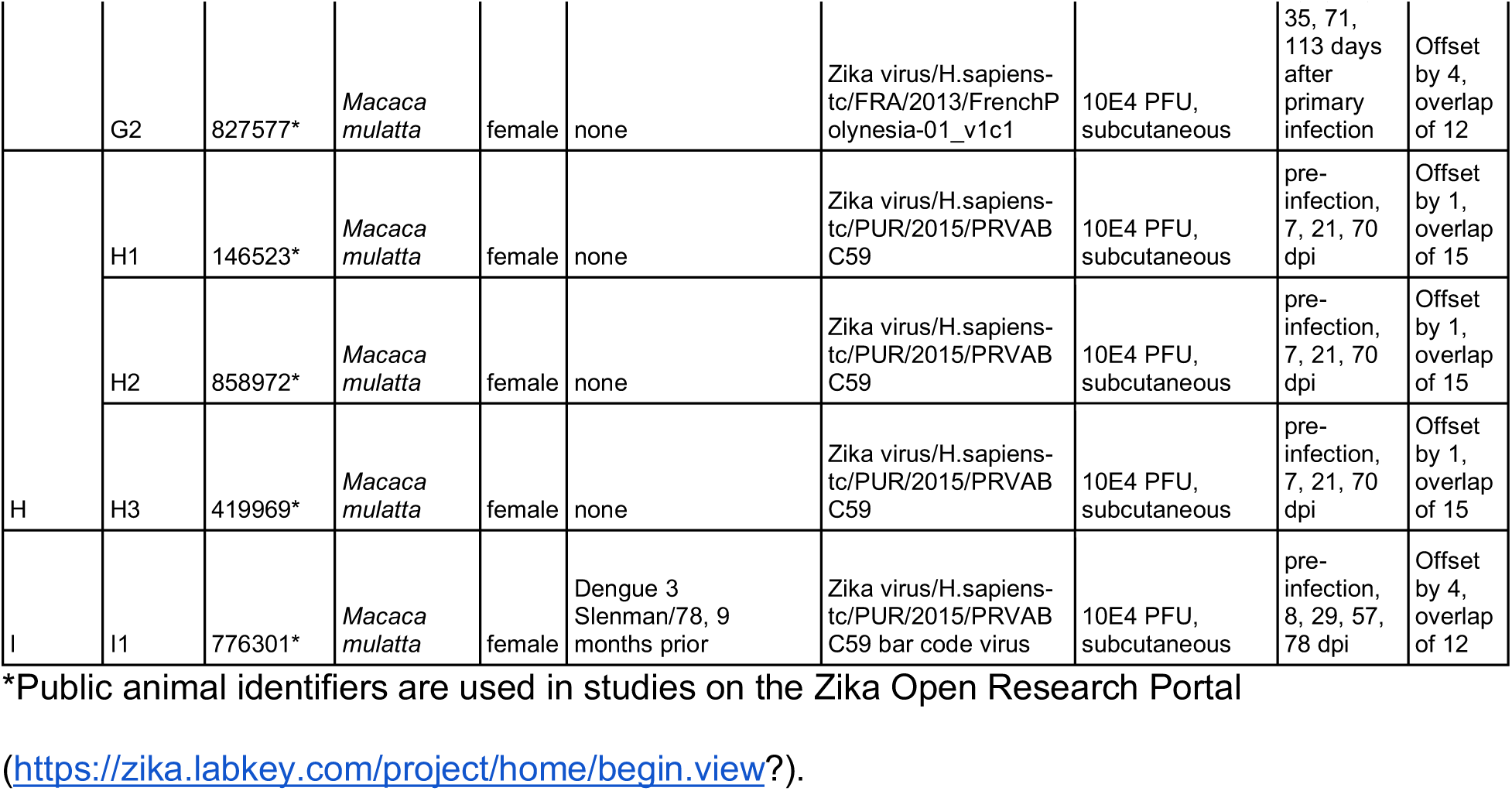
Animal demographics.

**Table 2.** Gestational details.

| Animal | Gestational days (gd) at inoculation | Pregnancy outcome |
| --- | --- | --- |
| G1 | 36 | delivery at 158 gd (122 dpi) |
| G2 | 38 | delivery at 158 gd (120 dpi) |
| H1 | 46 | fetectomy at 116 gd (70 dpi) |
| H2 | 47 | fetectomy at 117 gd (70 dpi) |
| H3 | 43 | fetectomy at 111 gd (68 dpi) |
| I1 | 35 | delivery at 155 gd (120 dpi) |

### Ethics

All monkeys were cared for by the staff at the Wisconsin National Primate Research Center (WNPRC) in accordance with the regulations and guidelines outlined in the Animal Welfare Act and the Guide for the Care and Use of Laboratory Animals and the recommendations of the Weatherall report (https://royalsociety.org/topics-policy/publications/2006/weatherall-report/). Per WNPRC standard operating procedures for animals assigned to protocols involving the experimental inoculation of an infectious pathogen, environmental enhancement included constant visual, auditory, and olfactory contact with conspecifics, the provision of feeding devices which inspire foraging behavior, the provision and rotation of novel manipulanda (e.g., Kong toys, nylabones, etc.), and enclosure furniture (i.e., perches, shelves). Per Animal Welfare Regulations (Title 9, Chapter 1, Subchapter A, Parts 1–4, Section 3.80 Primary enclosures) the animals were housed in a nonhuman primate Group 3 enclosure with at least 4.3 square feet of floor space and at least 30 inches of height. This study was approved by the University of Wisconsin-Madison Graduate School Institutional Animal Care and Use Committee (animal protocol numbers G005401 and G005443).

### Virus stocks

SIVmac239 (GenBank accession: M33262) stock was prepared by the WNPRC Virology Services Unit. Vero cells were transfected with the SIVmac239 plasmid. Infectious supernatant was then transferred onto rhesus PBMC. The stock was not passaged before collecting and freezing. ZIKV strain H/PF/2013 (GenBank accession: KJ776791) was obtained from Xavier de Lamballerie (European Virus Archive, Marseille, France) and passage history is described in Dudley et al. [16]. ZIKV strain MR766, ZIKV strain PRVABC59, and DENV-2 strain New Guinea C were generously provided by Brandy Russell (CDC, Ft. Collins, CO). ZIKV strain MR766 passage history has been described in Aliota et al. [38]. A molecularly-barcoded version of ZIKV strain PRVABC59 (Zika virus/H.sapiens-tc/PUR/2015/PRVABC59; GenBank accession: KU501215) was constructed as described in Aliota et al. [39]. DENV-2 strain New Guinea C (GenBank accession: FJ390389), originally isolated from a human in New Guinea, underwent 17 rounds of amplification on cells and/or suckling mice followed by a single round of amplification on C6/36 cells; virus stocks were prepared by inoculation onto a confluent monolayer of C6/36 mosquito cells. DENV-3 strain Sleman/78 was obtained from the NIH; virus stocks were prepared by a single passage on C6/36 cells.

### Multiple sequence alignment

Full-length ZIKV and DENV polyprotein sequences were extracted from the National Center for Biotechnology Information (NCBI) database into Geneious Pro 9.1.8 (Biomatters, Ltd., Auckland, New Zealand). These sequences included the ZIKV and DENV polyprotein sequences described above (see “Virus stocks”), as well as DENV-1 strain VR-1254 (GenBank accession: EU848545) and DENV-4 strain VID-V2055 (GenBank accession: KF955510). These amino acid sequences were aligned using the Geneious alignment algorithm as implemented in Geneious Pro 9.1.8 using default parameters (global alignment with free end gaps, cost matrix: Blosum62).

### Peptide array design and synthesis

Viral protein sequences were selected and submitted to Roche Sequencing Solutions (Madison, WI) for development into a peptide microarray as part of an early access program. Sequences included three ZIKV polyproteins (an Asian strain, ZIKV-FP, GenBank accession: KJ776791.2; an African strain, ZIKV-MR766, GenBank accession: KU720415.1; and an American strain, ZIKV-PR, GenBank accession: KU501215.1), four DENV polyproteins (DENV-1, GenBank accession: EU848545.1; DENV-2, GenBank accession: KM204118.1; DENV-3, GenBank accession: M93130.1; and DENV-4, GenBank accession: KF955510.1), and one SIVmac239 env protein (GenBank accession: AAA47637.1), which were used for most analyses in this study. Sequences used for cumulative distribution function (CDF) plots (see Figure 3) and analysis include sequences for mosquito-borne viruses found in Africa and known to infect humans [40], as well as one Japanese encephalitis virus strain (GenBank accession: KX945367.1). Accession numbers used to represent each viral protein are listed in the supplemental material (Table S7). Proteins were tiled as non-redundant 16 amino acid peptides, overlapping by 12 or 15 amino acids. The array designs are publicly available at https://go.wisc.edu/b726s1.

The peptide sequences were synthesized *in situ* with a Roche Sequencing Solutions Maskless Array Synthesizer (MAS) by light-directed solid-phase peptide synthesis using an amino-functionalized support (Geiner Bio-One) coupled with a 6-aminohexanoic acid linker and amino acid derivatives carrying a photosensitive 2-(2-nitrophenyl) propyloxycarbonyl (NPPOC) protection group (Orgentis Chemicals). Unique peptides were synthesized in random positions on the array to minimize impact of positional bias. Each array is comprised of twelve subarrays, where each subarray can process one sample and each subarray contains up to 392,318 unique peptide sequences.

### Peptide array sample binding

Macaque serum samples were diluted 1:100 in binding buffer (0.01M Tris-Cl, pH 7.4, 1% alkali-soluble casein, 0.05% Tween-20). Diluted sample aliquots and binding buffer-only negative controls were bound to arrays overnight for 16-20 h at 4°C. After binding, the arrays were washed 3x in wash buffer (1x TBS, 0.05% Tween-20), 10 min per wash. Primary sample binding was detected via 8F1-biotin mouse anti-primate IgG (NIH Nonhuman Primate Reagent Resource) secondary antibody. The secondary antibody was diluted 1:10,000 (final concentration 0.1 ng/µl) in secondary binding buffer (1x TBS, 1% alkali-soluble casein, 0.05% Tween-20) and incubated with arrays for 3 h at room temperature, then washed 3x in wash buffer (10 min per wash) and 30 sec in reagent-grade water. The secondary antibody was labeled with Cy5-Streptavidin (GE Healthcare; 5 ng/µl in 0.5x TBS, 1% alkali-soluble casein, 0.05% Tween-20) for 1 h at room temperature, then the array was washed 2x for 1 min in 1x TBS, and washed once for 30 sec in reagent-grade water. Fluorescent signal of the secondary antibody was detected by scanning at 635 nm at 2 µm resolution and 25% gain, using an MS200 microarray scanner (Roche NimbleGen).

### Peptide array data processing

The datafiles and analysis code for figures are available from https://go.wisc.edu/b726s1. All figures use the log base 2 of the raw fluorescence signal intensity values. For each sample, each unique peptide was assayed and processed once; then results from peptides redundant to multiple proteomes (i.e. were present in more than one strain represented) were restored to each protein.

For cumulative distribution function (CDF) plots, fluorescence signal intensities were log base 2 transformed and background reactivity in the blank (binding buffer only) control sample was subtracted for each peptide. The fold change from 0 dpi was calculated by subtracting reactivity at 0 dpi from reactivity at 28 dpi. To reduce instrument-related variance, the data was then filtered by taking the minimum intensity of two consecutive peptides with 1 amino acid offset, thereby reducing peptide outliers by ensuring measured reactivity occurs in multiple consecutive peptides.

### Validation of peptide array findings

To confirm the validity of our findings from this recently-developed peptide microarray platform, we assessed its performance against the humoral response produced by infection with simian immunodeficiency virus (SIV), which has been well-characterized by conventional methods such as enzyme-linked immunosorbent assays (ELISAs), enzyme-linked immunosorbent spot assays (ELISPOTs), epitope-prediction methods, or other protein arrays. We synthesized linear 16-mer peptides, overlapping by 12 amino acids, representing the SIVmac239 envelope protein (env) and analyzed serum from two Mauritian cynomolgus macaques for SIV-specific IgG reactivity before and approximately 125 days after SIVmac239 infection (Table 1). (We have previously plotted data procured from peptide array assays of these samples [37] while investigating the antibody response to SIV in the context of simian pegivirus infection; here we show the same data using the updated, improved data processing pipeline described above). Post-infection samples showed fluorescence intensity as high as 1,000 times the intensity in pre-infection samples (Supplemental figure S1). Regions of higher-fold increases in fluorescence intensity corresponded to previously defined variable domains of env which are known antibody targets, the variable loop regions, as well as others corresponding to known epitopes [41,42]. Taken together, these results validate epitope definition on the peptide microarray platform and show this platform to be capable of high-resolution virus-specific IgG epitope identification using the analytic methods utilized here.

## Results

### Identification of linear B cell epitopes in the ZIKV polyprotein

We sought to determine antibody binding, or reactivity, to the ZIKV polyprotein following ZIKV infection. We tiled 16-residue (16-mer) peptides overlapping by 15 amino acids representing different ZIKV polyproteins and evaluated the antibody binding of serum samples from animals with recent ZIKV-FP (animals B1, D1, and D2) or ZIKV-MR766 (animal C1) infections (Table 1). Animals in both groups had demonstrated neutralizing antibody responses at 28 dpi, measured by 90% plaque reduction neutralization tests (PRNT_90_) as described previously [16,38,23]. Peptides were defined as reactive if the signal intensity was greater after infection with a cognate strain (e.g., increased signal intensity against ZIKV peptides in an animal infected with ZIKV) than it was before infection. Peptides were defined as cross-reactive if the signal intensity was greater after infection with a noncognate strain (e.g., increased signal intensity against ZIKV peptides in an animal infected with DENV). Statistical significance of the change in signal intensity versus no change was calculated using a two-tailed log-ratio t-test. True epitopes were expected to induce antibody binding to multiple peptides with overlapping sequences; thus regions were only considered epitopes when there was a statistically significant increase in intensity in a post-infection sample relative to intensity in a pre-infection sample in three or more consecutive peptides.

Reactivity was most commonly observed in three regions of the flavivirus polyprotein: envelope protein, NS2B, and nonstructural protein 3 (NS3); therefore, most of the analysis in this paper is limited to these regions. Antibody binding to peptides from the envelope protein was seen in all four animals at 28 dpi; antibody binding in the NS3 region was seen in three out of four animals (Figure 1; reactivity throughout the entire ZIKV polyprotein can be seen in supplemental figure S2). The four animals exhibited similar responses against an Asian ZIKV strain, ZIKV-FP (Figure 1), as against an African strain and an American strain (ZIKV-MR766 and ZIKV-PR respectively, supplemental figure S3).

**Figure 1.**
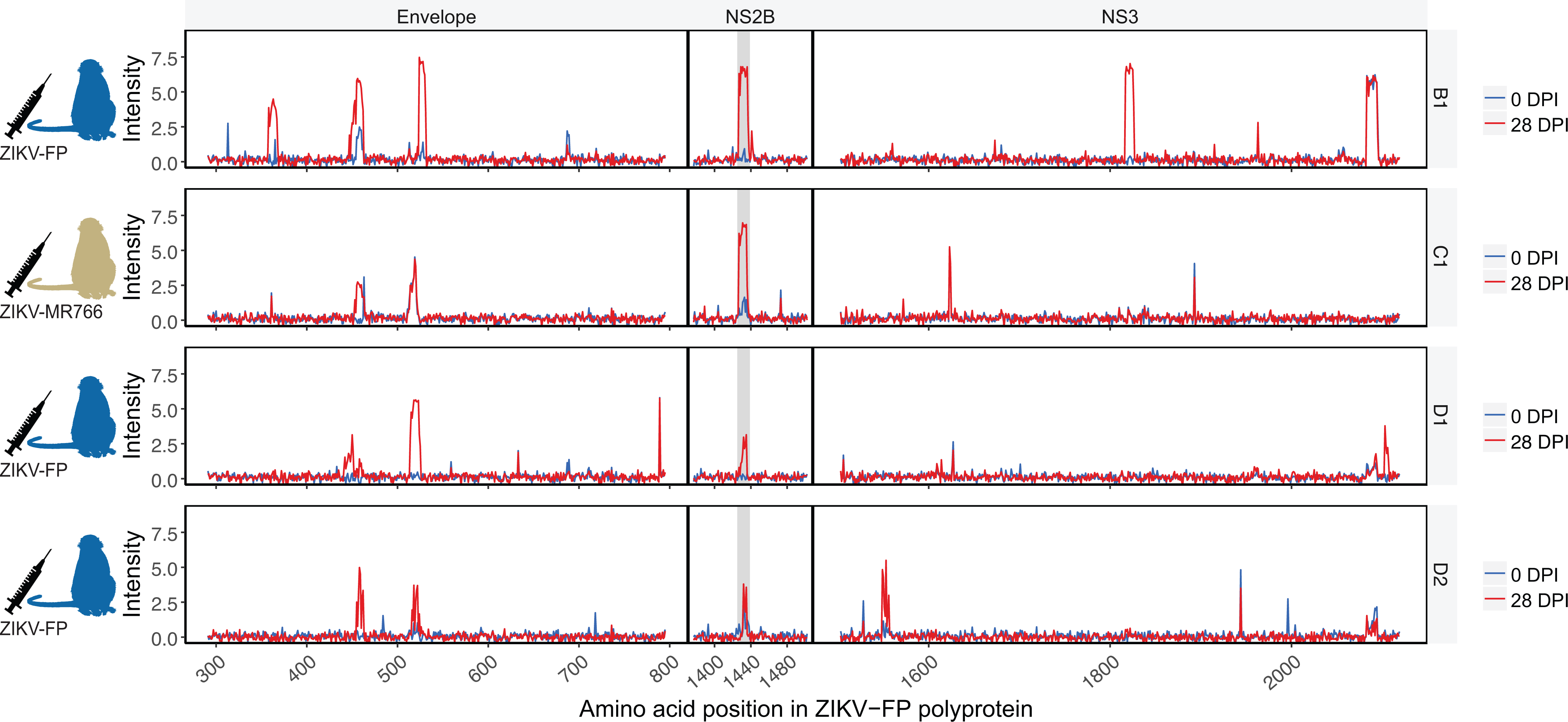
Reactivity of ZIKV-convalescent animals against the ZIKV-FP polyprotein. Serum from rhesus macaques infected with ZIKV-FP (animals B1, D1, and D2) and with ZIKV-MR766 (animal C1) was assayed for IgG recognition of the ZIKV-FP polyprotein. Reactivity prior to infection and at 28 dpi is shown. The NS2B_1427-1451_RD25 epitope is highlighted in grey.

All four animals exhibited antibody binding to the ZIKV NS2B epitope similar to that documented in humans [12]. Though other reactive epitopes were identified in other proteins in multiple animals, this epitope was the only epitope in our study for which all ZIKV-infected animals showed reactivity. All ten ZIKV-infected animals in this study were used to determine statistical significance of this epitope in order to avoid making statistical inferences using very small sample sizes [43]. Area under the curve (AUC) values and corresponding receiver operating characteristic (ROC) curves were calculated (supplemental figure S4) to identify ten peptides at positions 1427-1436 in the polyprotein (for a total of 25 amino acids, sequence RAGDITWEKDAEVTGNSPRLDVALD) as the best-performing epitope for which a statistically significant change from 0 dpi was observed, hereafter referred to as NS2B_1427-1451_RD25 (AUC of 0.9375, 95% confidence interval of 0.89065 to 0.9375). Using the mean signal intensity across the ten peptides, the change of signal intensity from 0 dpi to 21-28 dpi was statistically significant versus no change by a two-tailed log-ratio t-test (p-value = 0.000501 < 0.05, df = 9). This epitope is slightly longer than that found by Mishra et al. (nine 12-mer peptides, for a total of 20 amino acids, sequence DITWEKDAEVTGNSPRLDVA) [12].

### Cross-reactivity of ZIKV-convalescent serum with DENV polyproteins and with arbovirus proteomes

DENV polyproteins share an average of 55% sequence identity with ZIKV polyproteins [1,2], and the region of DENV NS2B corresponding to the immunoreactive ZIKV NS2B_1427-1451_RD25 epitope shares an average of 41% identity (Figure 2 A). To examine cross-reactivity, we analyzed the antibody binding of samples from the ZIKV-convalescent animals against DENV polyproteins represented on the array (Figure 2 B-E). Cross-reactivity was apparent, with antibodies from the ZIKV-convalescent/DENV-naive animals recognizing regions of DENV polyproteins. All four animals showed some cross-reactivity against the DENV envelope protein and DENV NS3. Animals exhibited comparable cross-reactivities to all four DENV serotypes, with the highest single instance of cross-reactivity observed against a DENV-3 epitope in the NS3 region. No significant cross-reactivity against the DENV NS2B protein was observed for these or any ZIKV-infected animals in this study (p-value = 0.548, df = 9).

**Figure 2.**
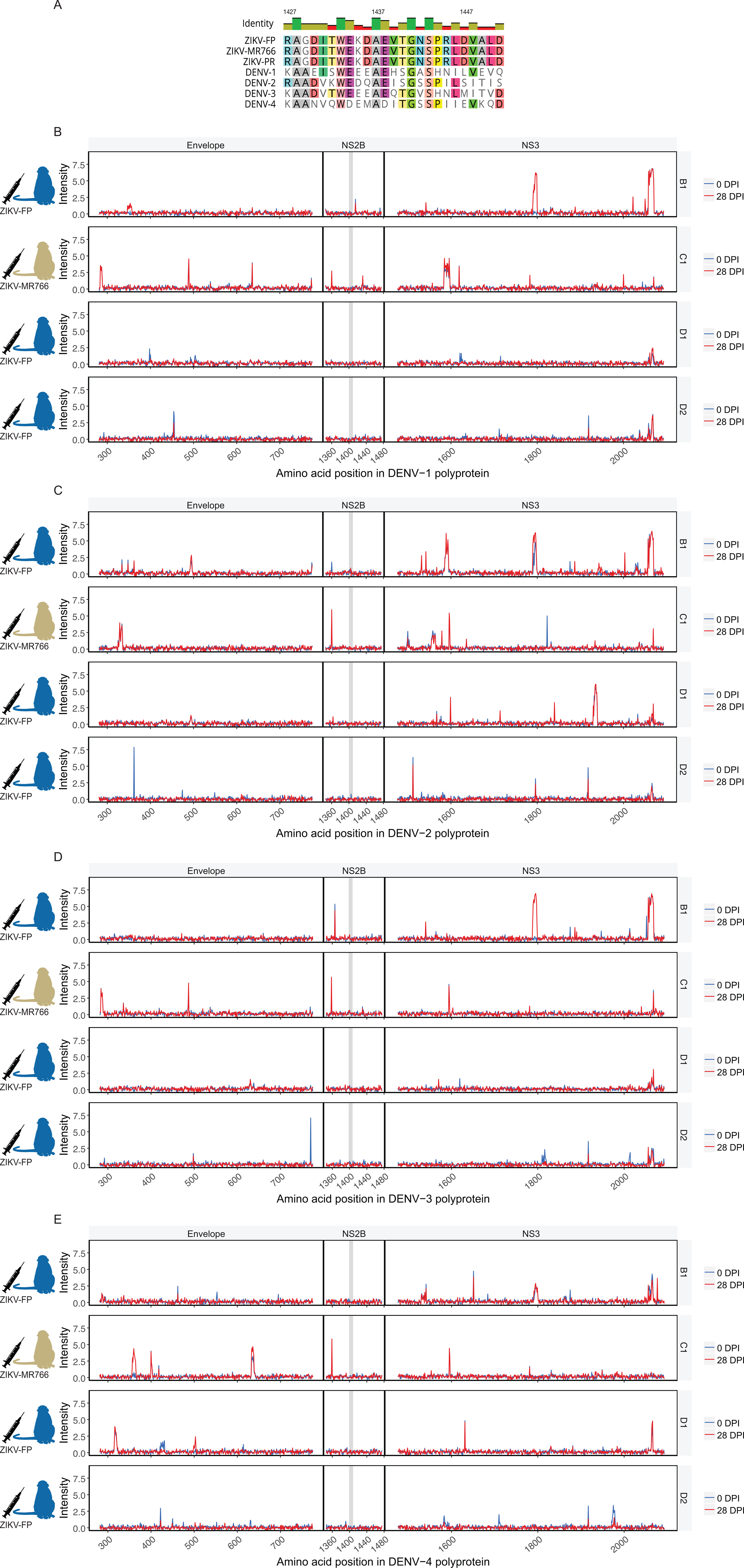
Cross-reactivity of ZIKV-convalescent animals against polyproteins of the four DENV serotypes. DENV serotypes share an average of 41% sequence similarity with the aligned ZIKV NS2B epitope (A). Serum from rhesus macaques infected with ZIKV-FP (animals B1, D1, and D2) or ZIKV-MR766 (C1) was assayed for cross-reactivity against polyproteins of DENV serotypes 1-4 (B-E, respectively). The segment of DENV NS2B which aligns with the ZIKV NS2B_1427-1451_RD25 epitope is highlighted in grey.

Given this array’s capacity to screen for antibody binding to peptides representing many different virus proteomes at once, we also assessed cross-reactivity against the 27 other arboviruses (for a total of 28 arboviruses) represented. We plotted the fold change in reactivity from 0 to 28 dpi for each peptide in each virus’s proteome using cumulative distribution function (CDF) plots (Figure 3). CDF plots were used to determine whether the sum of reactivity across a viral proteome could distinguish reactivity to the infecting pathogen (in this case, ZIKV) from cross-reactivity to a variety of other similar or dissimilar pathogens (in this case, 27 other arboviruses). Cross-reactivity in peptides in other arboviruses was observed, but it was rare compared with the reactivity seen in the ZIKV peptides. The greatest degree of cross-reactivity occurred among flaviviruses, in particular Ntaya virus, Spondweni virus, Bagaza virus, and DENV-3 (Figure 3 C). Pooled t-tests showed significant differences between reactivity against ZIKV strains and other viruses for each of the four animals (Table 3). In animal B1, reactivity against ZIKV was significantly different from cross-reactivity against all other arboviruses (p-values ranging from <0.0001 to 0.0006). Animal C1’s reactivity against ZIKV was significantly different (p-values <0.05) for all viruses assayed except Babanki virus, Banzi virus, DENV-4, Uganda S virus, and yellow fever virus. D1 showed reactivity against ZIKV that was significantly different from reactivity against all other viruses except Spondweni virus. D2 showed more significant cross-reactivity with other arboviruses; in D2, ZIKV reactivity differed significantly from reactivity to all other viruses except Bwamba virus, Chikungunya virus, DENV-1, Middelburg virus, Ndumu virus, Rift Valley fever virus, Spondweni virus, and Uganda S virus. Of note, D2 did also exhibit the smallest increase in fold change from 0 to 28 dpi in reactivity against ZIKV; this higher likelihood of cross-reactivity may be attributable to the minimal overall amount of reactivity present. Reactivity for the two ZIKV strains compared, one African strain (ZIKV-MR766, GenBank accession: NC_012532.1) and one Asian/American strain (GenBank accession: NC_035889.1) was not significantly different for any animal (p-values ranging from 0.62 to 0.88).

**Figure 3.**
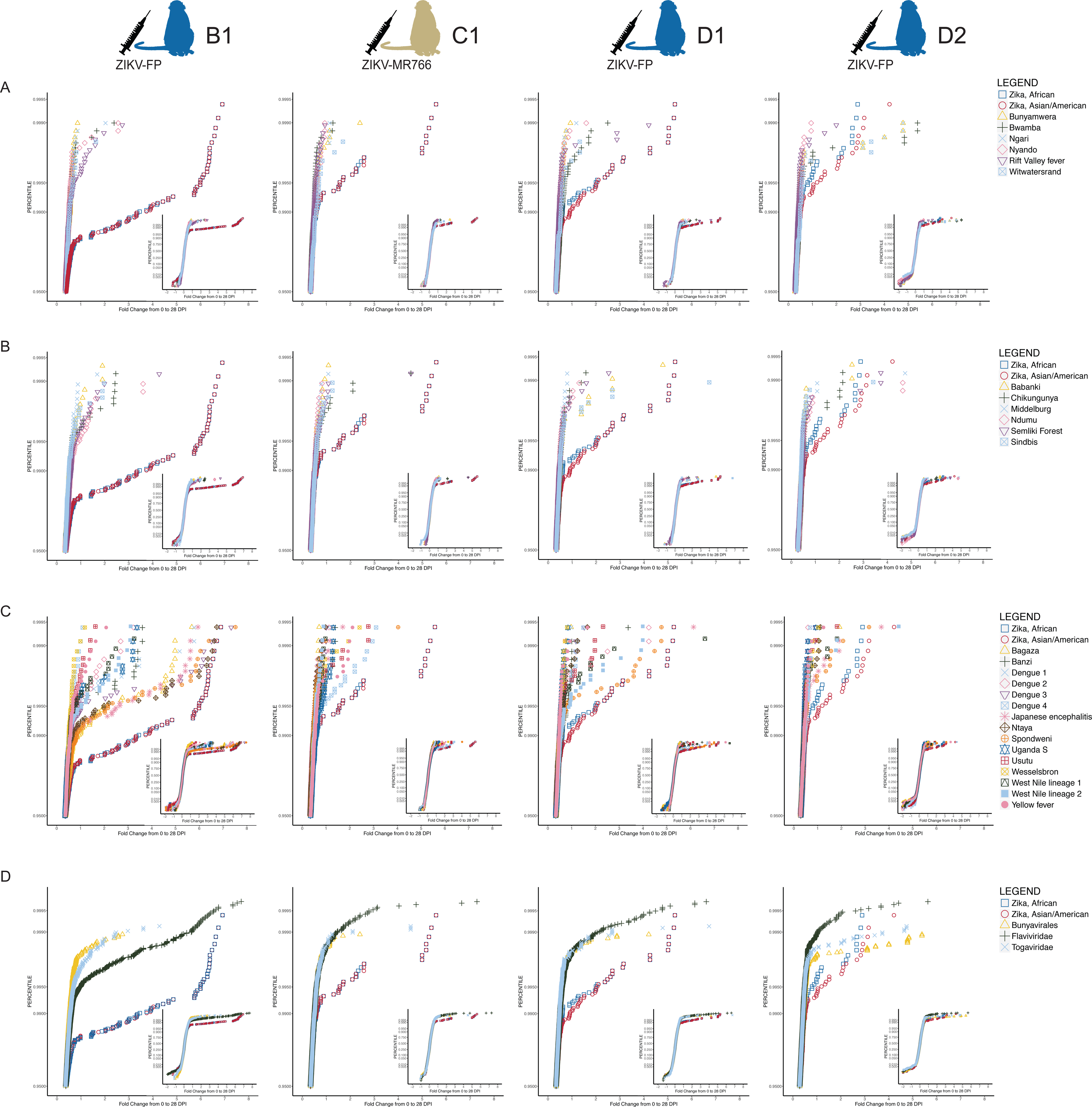
Cross-reactivity of sera from ZIKV-convalescent animals against the complete polyproteins or proteomes of 27 arboviruses represented on the array. Cumulative distribution function (CDF) plots of fold change from 0 dpi to 28 dpi by of animals’ reactivity and cross-reactivity to different viral proteomes are shown; data for animals infected with ZIKV-FP (B1, D1, and D2) and ZIKV-MR766 (C1) are shown. Reactivity to two ZIKV strains, one African and one Asian/American, is compared against all viruses on the array of order Bunyavirales (A), of family Togaviridae (B), of family Flaviviridae (C), and with the averages of reactivity of all viruses in Bunyavirales, of all viruses in Togaviridae, and of all viruses in Flaviviridae (D). The region of interest is shown large in the figure, while the full CDF plot is shown as an inset. ZIKV strains demonstrate significantly increased fold change in reactivity compared to the majority of other viruses represented on the array (see Table 3).

**Table 3.** Significance of difference from ZIKV of summed reactivity.

| Arbovirus species | p-value for t-test against Asian/American ZIKV strain |  |  |  |
| --- | --- | --- | --- | --- |
|  | B1 | C1 | D1 | D2 |
| Babanki | <0.0001 | 0.0924* | 0.0188 | 0.0004 |
| Bagaza | <0.0001 | 0.0010 | 0.0001 | <0.0001 |
| Banzi | <0.0001 | 0.0609* | 0.0060 | 0.0062 |
| Bunyamwera | <0.0001 | 0.0094 | 0.0001 | 0.0202 |
| Bwamba | <0.0001 | 0.0019 | 0.0030 | 0.2367* |
| Chikungunya | <0.0001 | 0.0452 | 0.0002 | 0.0562* |
| Dengue 1 | <0.0001 | 0.0010 | 0.0015 | 0.2522* |
| Dengue 2 | <0.0001 | 0.0006 | 0.0053 | 0.0074 |
| Dengue 3 | 0.0001 | 0.0067 | 0.0017 | <0.0001 |
| Dengue 4 | <0.0001 | 0.1880* | <0.0001 | 0.0038 |
| Japanese encephalitis | 0.0002 | 0.0349 | 0.0045 | 0.0447 |
| Middelburg | <0.0001 | 0.0081 | 0.0006 | 0.2074* |
| Ndumu | <0.0001 | 0.0008 | <0.0001 | 0.1681* |
| Ngari | <0.0001 | 0.0110 | 0.0015 | 0.0997* |
| Ntaya | 0.0002 | 0.0002 | <0.0001 | 0.0476 |
| Nyando | <0.0001 | 0.0027 | <0.0001 | 0.0001 |
| Rift Valley fever | <0.0001 | 0.0085 | 0.0001 | 0.1868* |
| Semliki Forest | <0.0001 | 0.0118 | <0.0001 | 0.0053 |
| Sindbis | <0.0001 | 0.0417 | 0.0010 | <0.0001 |
| Spondweni | 0.0006 | 0.0015 | 0.5734* | 0.2896* |
| Uganda S | <0.0001 | 0.0608* | <0.0001 | 0.2134* |
| Usutu | <0.0001 | 0.0129 | 0.0084 | 0.0053 |
| Wesselsbron | <0.0001 | 0.0032 | 0.0005 | 0.0002 |
| West Nile lineage 1 | <0.0001 | 0.0007 | 0.0035 | <0.0001 |
| West Nile lineage 2 | <0.0001 | 0.0009 | 0.0052 | 0.0016 |
| Witwatersrand | <0.0001 | 0.0083 | 0.0034 | 0.0289 |
| Yellow fever | <0.0001 | 0.5379* | 0.0003 | 0.0002 |
| Zika, African | 0.7952* | 0.6184* | 0.8688* | 0.7304* |

\*Non-significant values (p-value > 0.05).

### Anti-ZIKV antibody response during pregnancy

Given the importance of determining an accurate ZIKV infection history in pregnancy, we sought to characterize the gestational anti-ZIKV antibody response. We used the peptide microarray to evaluate serum samples from six pregnant animals. Animals were inoculated with ZIKV at 35-47 days post-conception (gestational date, gd) (Table 2). Two animals (G1 and G2) had no history of flavivirus exposure and were inoculated with ZIKV-FP at 36-38 gd. Three more flavivirus-naive animals (H1, H2, and H3) were infected with ZIKV-PR at 45-47 gd. One animal (I1) had a history of exposure to DENV-3 nine months prior to inoculation with a barcoded clone of ZIKV-PR at 35 gd [39]. Serum samples collected approximately one week post-infection and at two to six week intervals thereafter were analyzed against ZIKV polyproteins represented on the peptide array (Figure 4). All six pregnant animals exhibited anti-NS2B_1427-1451_RD25 reactivity by the early convalescent phase (21-29 dpi), though time of initial appearance and duration of the response varied between animals.

**Figure 4.**
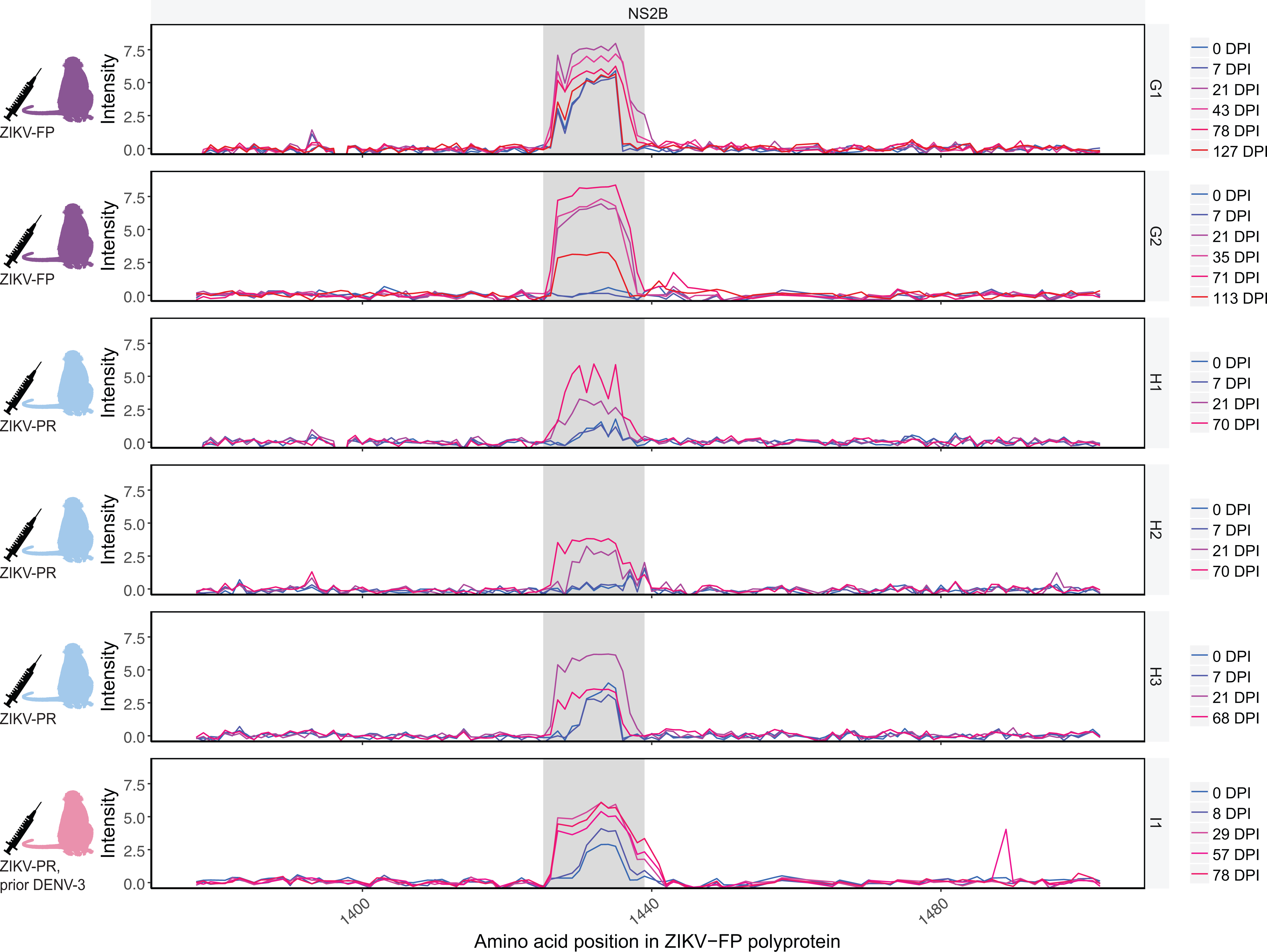
IgG reactivity against ZIKV NS2B_1427-1451_RD25 during pregnancy. Serum samples taken throughout the course of six animals’ pregnancies were evaluated against the ZIKV-FP polyprotein represented on peptide microarrays. Two animals (G1 and G2) were infected with ZIKV-FP at 36-38 gd. Three animals (H1, H2, H3) were infected with ZIKV-PR at 45-47 gd. One animal (I1) had been infected with DENV-3 nine months prior and was infected with a barcoded clone of ZIKV-PR at 35 gd. The NS2B_1427-1451_RD25 epitope is highlighted in grey. Reactivity against the ZIKV-FP envelope and NS3 proteins can be found in supplemental figure S5.

All pregnant animals showed a similar pattern of anti-ZIKV reactivity, with some differences in time to peak reactivity and duration of detectable reactivity. In G1, elevated baseline intensity in the ZIKV NS2B_1427-1451_RD25 region was present prior to inoculation with ZIKV-FP. Reactivity peaked in the acute phase (7 dpi) and subsequently decreased but remained above pre-infection levels through 127 dpi, after G1 had given birth. In G2, reactivity was not appreciable until the early convalescent phase (21 dpi) and peaked at 35 dpi, remaining elevated relative to pre-infection levels through the latest time point analyzed (113 dpi). Animals H1 and H3 showed elevated pre-infection intensity against NS2B_1427-1451_RD25. All three animals in cohort H showed reactivity by 21 dpi. H1 and H2 continued to exhibit increased reactivity through the latest time point measured (70 dpi), while H3 had peak reactivity at 21 dpi and decreased after. I1, an animal nine months post-DENV-3 infection, showed a pattern of reactivity similar to that in other animals, with anti-NS2B_1427-1451_RD25 IgG reactivity first appearing at 8 dpi and peaking at 29 dpi at a level 3.6 times pre-infection reactivity (Figure 4). Reactivity remained close to peak reactivity through 78 dpi.

All pregnant animals’ cross-reactivity against DENV polyproteins mirrored that seen in non-pregnant animals (supplemental figure S6).

### Differentiating DENV serology in ZIKV-immune animals

Previous assays have struggled to distinguish DENV serologic responses from ZIKV serologic responses [5–8]. We investigated whether the peptide microarray technology, and specifically reactivity patterns using the ZIKV NS2B_1427-1451_RD25 epitope, could distinguish DENV from ZIKV infections. Three animals (cohort F) had been challenged twice with ZIKV-FP 12 months prior and 9.5 months prior [16]; we collected serum from these animals, infected them with DENV-2, and collected serum 28 days after. These samples were analyzed against 16-mer peptides, with amino acid overlap of 12, representing DENV-2 and ZIKV-FP (Figure 5).

**Figure 5.**
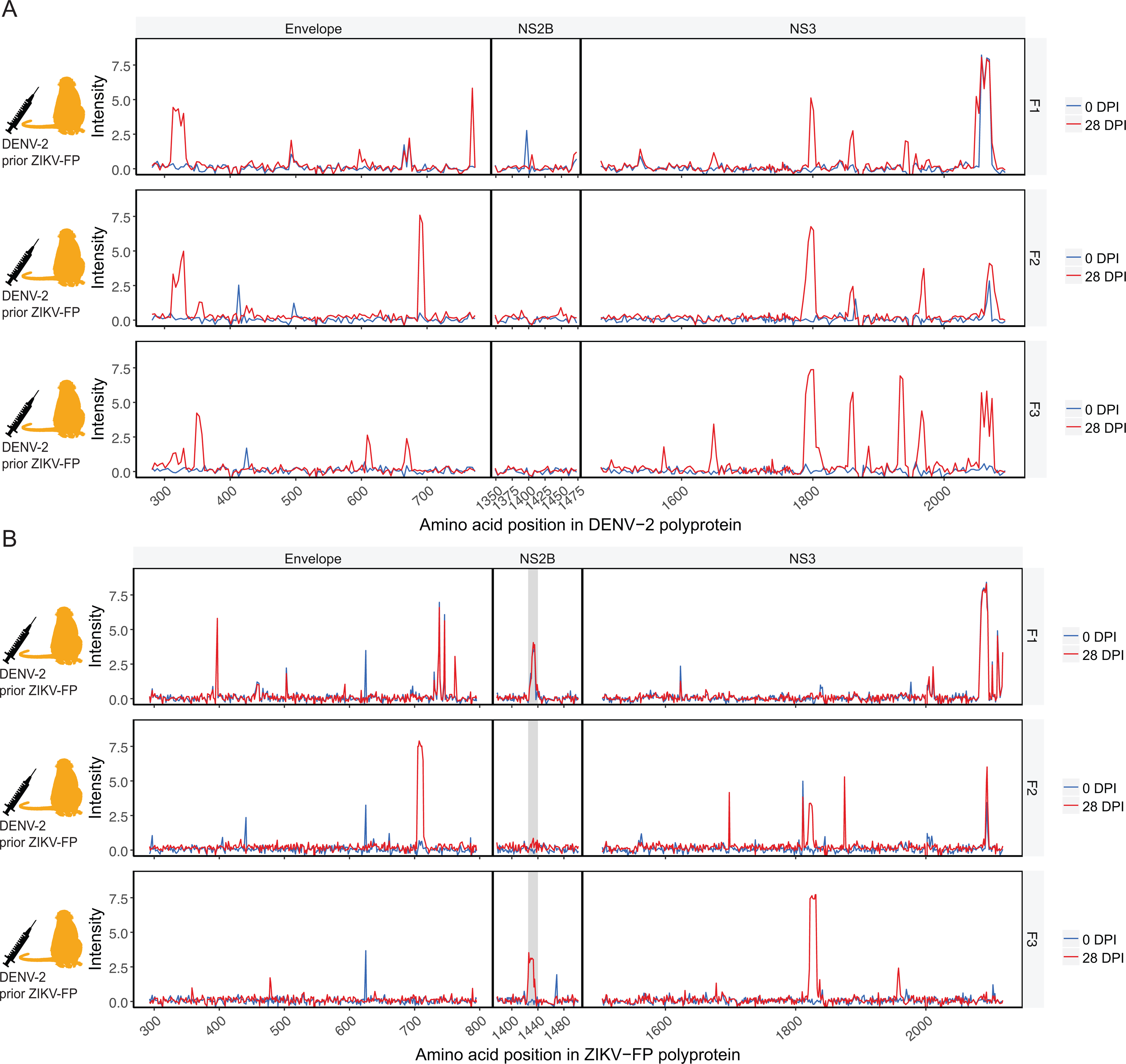
Antibody reactivity to DENV and ZIKV polyproteins of animals with recent DENV infection. Three macaques (F1, F2, and F3), with history of challenge and rechallenge with ZIKV-FP 12 and 9.5 months prior, were infected with DENV-2 and serum samples taken before DENV infection and at 28 dpi were assessed. Reactivity of these animals against peptides representing the envelope, NS2B, and NS3 proteins of DENV-2 (A) and ZIKV-FP (B) is shown. The ZIKV NS2B_1427-1451_RD25 epitope (in B) is highlighted in grey.

These animals showed reactivity against all four DENV polyproteins in regions representing the DENV envelope protein, DENV NS3, and others (Figure 5A) and cross-reactivity against corresponding regions of the ZIKV polyprotein (Figure 5B). One animal (F3) out of the three showed significant reactivity to the NS2B_1427-1451_RD25 epitope, though this reactivity was slightly outside the area of typical NS2B_1427-1451_RD25 reactivity (peptides 1421-1444 rather than 1427-1451). Another (F1) showed elevated pre-DENV infection intensity in NS2B_1427-1451_RD25 that did not change following DENV infection, possibly as a result of the animal’s prior ZIKV exposure.

Animal F2 showed no detectable NS2B_1427-1451_RD25 reactivity before or after exposure.

## Discussion

We describe the antibody binding of the anti-ZIKV IgG response in non-pregnant and pregnant rhesus macaques and compare this to the anti-DENV response. Using a recently developed high-density peptide microarray we show that macaques infected with ZIKV produce IgG antibodies which bind throughout the ZIKV polyprotein, including conserved antibody binding to an epitope in ZIKV NS2B, NS2B_1427-1451_RD25, which is apparent regardless of the ZIKV strain used for infection. We establish that cross-reactivity exists between anti-ZIKV and anti-DENV antibodies for the ZIKV and DENV polyproteins, and we show this technology can be used to differentiate anti-ZIKV reactivity from cross-reactivity to many other arboviruses. Additionally, we show the anti-NS2B_1427-1451_RD25 IgG response is susceptible to false positives in the context of DENV infection and may be susceptible to false-positives in flavivirus-immune individuals. Thus, while this epitope may be broadly useful for serosurveillance, it should be used with caution in the diagnosis of individual infections.

As has been seen in previous assays [2,12,44-46], we observed antibody cross-reactivity between ZIKV- and DENV-immune sera, though reactivity to the ZIKV NS2B_1427-1451_RD25 epitope was observed in all cases of recent ZIKV infection (10 out of 10 ZIKV-infected animals), and in only one out of three cases of recent DENV infection in animals with history of ZIKV exposure. Reactivity to the NS2B epitope was conserved whether the infecting strain was African, Asian, or American in origin. All ZIKV-infected animals produced anti-ZIKV IgG against the NS2B_1427-1451_RD25 epitope, though reactivity was sometimes small in magnitude (as in animal D2) or was measurable for only a short duration (as in animal H3). The lack of uniformity of the anti-ZIKV IgG response is especially relevant since all ZIKV-exposed animals could be followed, in contrast to studies in humans where there may be a selection bias for individuals with symptomatic ZIKV, which is thought to account for only approximately half of ZIKV infections [12,47]. Though anti-NS2B_1427-1451_RD25 reactivity was consistently detectable in early convalescence, it decayed in all but one case, that of a pregnant animal with previous DENV exposure, during the time period assessed. These findings corroborate findings in humans, in which symptomatic ZIKV infection was strongly associated with detectable anti-NS2B antibodies in the early convalescent phase (96%), but was less likely six months post-infection (44%) [12]. These results demonstrate the need for further investigation into the longevity and kinetics of the anti-ZIKV humoral response. Given that no other purported ZIKV-specific epitopes have been identified to date, these results also call into question how useful current serological methods may be in differentiating past ZIKV exposure from exposure to other flaviviruses. Additionally, one animal (F3) with a history of previous ZIKV infections showed reactivity against the ZIKV NS2B_1427-1451_RD25 epitope following experimental DENV infection, indicating that a history of ZIKV infection or a recent DENV infection has potential to confound results using this epitope. Thus, while this epitope appears to be more ZIKV-specific than most, its utility in guiding development of diagnostics will likely be limited.

The ability of the peptide array to differentiate seroreactivity against ZIKV from cross-reactivity to other viruses and to show cross-reactivity to 27 other mosquito-borne arboviruses suggests this technology, with additional optimization, could be useful for determining the etiology of fever of unknown origin and other non-specific symptoms in areas where mosquito-borne diseases are common. All ZIKV-infected animals showed the greatest fold change in reactivity against ZIKV proteomes. Though these differences in fold change did not always reach significance for all strains in all animals, the tendency to show the greatest reactivity in ZIKV strains, even in comparisons against very closely-related flaviviruses, for ZIKV-infected animals demonstrates the potential of this technology in guiding diagnostic development in the future.

This assay detected anti-NS2B_1427-1451_RD25 reactivity in all pregnant macaques exposed to ZIKV. Production of IgG antibodies has relevance to both mother and fetus, since maternal IgG crosses the placenta and transport of IgG across the placental barrier increases throughout the course of pregnancy [48]. It can be presumed the anti-ZIKV IgG produced by these animals also reached their fetuses, but whether these anti-NS2B_1427-1451_RD25 antibodies provide protection, contribute to the pathogenesis of ZIKV disease and ZIKV congenital effects, or are irrelevant in ZIKV pathology is currently unknown. Though we did not note any variation in outcomes with differences in antibody responses, it is possible deviations in antibody responses during pregnancy could help explain differences in outcomes following gestational ZIKV infection.

Several pregnant animals also showed elevated pre-infection intensity at the NS2B_1427-1451_RD25 epitope. This phenomenon may be due to innate immunodominance of the NS2B_1427-1451_RD25 epitope, or it may be due to molecular mimicry, though we have not found this epitope present in any other pathogen. These findings merit further and more thorough investigation than this current study can provide.

The peptide array technology used in this study has several limitations. The assay’s utility could be increased by the addition of quantitative capacities. We currently use a 1:100 antibody dilution since we have found this to produce an optimal signal:noise ratio, but it is possible serial dilutions could allow for measurement of quantitative results. Greater confidence in this assay’s results could be derived from assessing its ability to identify the binding of well-characterized monoclonal antibodies. Once validated in this way, the array could then be used to determine the specificity of new monoclonal antibodies as they are discovered. Additionally, our study defined a positive response as an increase from an animal’s pre-infection intensity; human patients usually cannot give pre-infection samples and must rely upon controls determined from humans having no known history of exposure to certain pathogens. The development of such a control would raise the specificity of the assay at the expense of sensitivity, and thus would risk missing some true positive results when definitive pre-infection samples from the same individual are not available.

The peptide array approach is also limited due to its reliance on continuous linear epitopes. Many documented ZIKV epitopes are conformational discontinuous epitopes [49–55]. Other epitope discovery methods will likely remain superior in defining discontinuous epitopes, but this technology is useful in identifying immunoreactive regions not previously considered as potential epitopes. Past work from our laboratory, including results from some of the animals whose sera was analyzed here, has shown anti-ZIKV neutralizing antibody titers measured by PRNT_90_ do not drop off but remain elevated as late as 64 dpi [16]. The discordance between levels of anti-NS2B_1427-1451_RD25 IgG and neutralizing antibody titers may indicate the anti-NS2B_1427-1451_RD25 antibodies do not play a role in protection against future infections, which may explain the drop-off in their production observed in macaques and in humans.

In the future, this technology could be expanded for use in profiling antibody responses to many other pathogens. This tool was able to detect known and previously unknown epitopes throughout the ZIKV proteome, including epitopes in unexpected regions such as NS2B. This approach could be applied to other NTDs to advance diagnostic and vaccine development. The array used in this study simultaneously evaluates antibody responses against the entire proteomes of every mosquito-borne virus known to infect humans in Africa. Building off what we have learned through this and other peptide array analyses, we intend to use this array to survey antibody reactivity in African populations, through which we may identify previously unknown epitopes for some of the rare pathogens represented on the array. Additionally, we plan to use this assay to evaluate and compare both IgM and IgG responses in these analyses, which may help elucidate the kinetics of the immediate and long-term antibody response to pathogens.

In summary, this work in macaques demonstrates the capacity of a recently developed peptide microarray to profile the binding of distinct anti-ZIKV and anti-DENV IgG antibody responses in experimental infections. Our work shows the anti-NS2B_1427-1451_RD25 IgG response is characterized by relatively rapid decay and is susceptible to confounding, mirroring results seen in humans. The peptide microarray technology used shows particular promise in evaluating full-proteome antibody binding for a large number of pathogens efficiently and may be especially useful for neglected tropical diseases for which diagnostics are rudimentary or non-existent.

## Acknowledgments

We would like to thank Adam Ericsen for helpful discussions.

Grant support: National Institutes of Health grants R01AI116382 and R24OD017850 (ASH, AKH, CRB, AB, DMD, CMN, MK, MEB, MR, LMS, DHO) and the Pediatric Infectious Diseases Society Fellowship Award funded by Stanley A. Plotkin and Sanofi Pasteur (ELM).

## Supplemental material

**Supplemental figure S1**. SIV env epitopes identified by a high-density peptide microarray overlap known epitopes in variable loop regions. Serum from two Mauritian cynomolgus macaques (animals A1 and A2) before and after infection with SIVmac239 was evaluated for reactivity to overlapping peptides representing the SIVmac239 env protein sequence on a linear peptide microarray. SIV env variable loop regions [41] are highlighted in grey.

**Supplemental figure S2**. Reactivity of B1, C1, D1, and D2 against the complete polyprotein of ZIKV-FP. The NS2B_1427-1451_RD25 epitope is highlighted in grey.

**Supplemental figure S3**. Reactivity of B1, C1, D1, and D2 against ZIKV-MR766 (A) and ZIKV-PR (B). The NS2B_1427-1451_RD25 epitope is highlighted in grey.

**Supplemental figure S4**. ROC curves for best-performing epitope and multidimensional scaling (MDS) plot for different time points. ROC curves (A) were generated using data from the 8 animals analyzed against ZIKV polyproteins tiled as 16 amino acid peptides overlapping by 15 amino acids since this was the most complete dataset. ROC curves were created using the 8 samples collected at 0 DPI as the control group and the same 8 samples collected at 21-28 DPI samples as the test group. We chose consecutive peptides within the NS2B region that maximized the differences of mean log-normalized intensities between the controls and test samples. The decision threshold was determined by maximizing the AUC and the resulting ROCs of 9 to 13 peptides were plotted. ROC curves overlapped. MDS plots (B) of distances between the log-normalized gene expression profiles were created using Limma plot MDS R library. Only the 10 peptides in the identified NS2B_1427-1451_RD25 epitope were used. Based on MDS analysis, pre-infection samples (blue), samples from early convalescence (21-28 dpi, red), and samples from any later time (>43 dpi, yellow) clustered in separate groups.

**Supplemental figure S5.** Reactivity of cohort G (ZIKV-FP infection during first trimester), H (ZIKV-PR infection during first trimester), and I (history of DENV-3 infection, ZIKV-PR infection during first trimester) against the ZIKV-FP polyprotein. The NS2B_1427-1451_RD25 epitope is highlighted in grey.

**Supplemental figure S6.** Reactivity of cohort G (ZIKV-FP infection during first trimester), H (ZIKV-PR infection during first trimester), and I (history of DENV-3 infection, ZIKV-PR infection during first trimester) against the DENV-2 polyprotein. The region of the DENV-2 polyprotein corresponding to the ZIKV NS2B_1427-1451_RD25 epitope is highlighted in grey.

**Supplemental Table S7.** Arboviral proteins and polyproteins used for CDF plot analysis.

